# Transcriptomic analysis of the lesser spotted catshark (*Scyliorhinus canicula*) pancreas, liver and brain reveals molecular level conservation of vertebrate pancreas function

**DOI:** 10.1101/006056

**Authors:** John F Mulley, Adam D Hargreaves, Matthew J Hegarty, R. Scott Heller, Martin T Swain

## Abstract

**Background:** Understanding the evolution of the vertebrate pancreas is key to understanding its functions. The chondrichthyes (cartilaginous fish such as sharks and rays) have been suggested to possess the most ancient example of a distinct pancreas with both hormonal (endocrine) and digestive (exocrine) roles, although the lack of genetic, genomic and transcriptomic data for cartilaginous fish has hindered a more thorough understanding of the molecular-level functions of the chondrichthyan pancreas, particularly with respect to their “unusual” energy metabolism (where ketone bodies and amino acids are the main oxidative fuel source) and their paradoxical ability to both maintain stable blood glucose levels and tolerate extensive periods of hypoglycemia. In order to shed light on some of these processes we have carried out the first large-scale comparative transcriptomic survey of multiple cartilaginous fish tissues: the pancreas, brain and liver of the lesser spotted catshark, *Scyliorhinus canicula*.

**Results:** We generated a mutli-tissue assembly comprising 86,006 contigs, of which 44,794 were assigned to a particular tissue or combination of tissue based on mapping of sequencing reads. We have characterised transcripts encoding genes involved in insulin regulation, glucose sensing, transcriptional regulation, signaling and digestion, as well as many peptide hormone precursors and their receptors for the first time. Comparisons to published mammalian pancreas transcriptomes reveals that mechanisms of glucose sensing and insulin regulation used to establish and maintain a stable internal environment are conserved across jawed vertebrates and likely pre-date the vertebrate radiation. Conservation of pancreatic hormones and genes encoding digestive proteins support the single, early evolution of a distinct pancreatic gland with endocrine and exocrine functions in vertebrates, although the peptide diversity of the early vertebrate pancreas has been overestimated as a result of the use of cross-reacting antisera in earlier studies. A three hormone islet organ is therefore the basal vertebrate condition, later elaborated upon only in the tetrapod lineage.

**Conclusions:** The cartilaginous fish are a great untapped resource for the reconstruction of patterns and processes of vertebrate evolution and new approaches such as those described in this paper will greatly facilitate their incorporation into the rank of “model organism”.

## Background

Chondrichthyans (cartilaginous fish such as sharks, skates, rays (elasmobranchs) and chimeras (holocephalans)) possess the earliest example of a distinct pancreatic gland containing multiple cell types with both endocrine and exocrine functions in vertebrates [1, 2]. The more basal (“primitive”) vertebrate lineages such as the jawless hagfish and lampreys (Figure 1) possess only small islet organs containing insulin- and somatostatin-producing endocrine cells and these islets lack any glucagon-producing cells or exocrine function [2–4]. The accumulation of multiple cell types into a single compact gland was an important step in pancreas evolution (and can be considered to be a vertebrate innovation) [5, 6] and it has been suggested that a switch from sensing gut-glucose to blood-glucose to establish a “stable inner *milieu*” via homeostatic mechanisms may have been an important factor in the evolution of a more complex glucose-dependent brain in vertebrates, protected from hyper- and hypoglycaemia [7–9]. However, the fact that insulin-like peptides in insects seem to fulfil similar roles in glucose metabolism and other physiological processes such as growth and reproduction suggests that at least elements of these mechanisms may have a more ancient origin [10].

**Figure 1.**
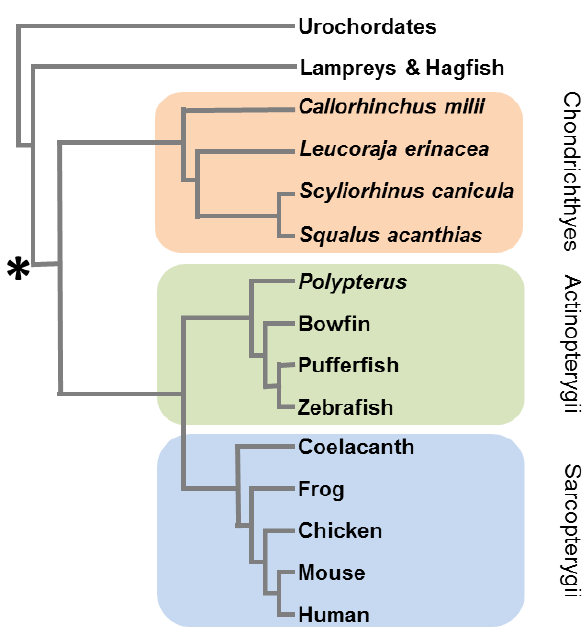
Phylogenetic tree of the major extant vertebrate groups. The relationships of the most common chondrichthyan (cartilaginous fish) model species (Elephant shark, *Callorhinchus milii*; Little skate, *Leucoraja erinacea*; Lesser spotted catshark, *Scyliorhinus canicula*; Spiny dogfish, *Squalus acanthias*) are shown, as are representative lineages from the ray-finned (actinopterygian) and lobe-finned (sarcopterygian) fish. The origin of the combined endocrine and exocrine pancreatic gland at the base of the jawed vertebrates is indicated.

Decades of research using light and electron microscopy and immunohistochemistry has revealed a great deal about the structure and organisation of the chondrichthyan pancreas and more recent studies have characterised the protein sequence and structure of some of the key pancreatic hormones [11–15]. The endocrine islets of chondrichthyans are typically scattered within exocrine tissue, sometimes associated with minor ducts and contain a number of distinct cell types, thought to include the four major pancreas cell types: α-cells producing glucagon to increase blood glucose; β-cells producing insulin to reduce blood glucose; δ-cells producing somatostatin to regulate pancreatic hormones such as insulin and glucagon and γ-cells producing pancreatic polypeptide [16–18]. In addition to structural, cellular and hormonal conservation of the chondrichthyan pancreas compared to other vertebrates, there is also a conservation of function, with glucose-sensitive insulin release [11], pancreatectomy-induced hyperglycemia [19, 20] and exogenous insulin-induced hypoglycaemic effects [11, 21–23]. However, although blood plasma glucose levels are maintained at a fairly constant level during feeding and fasting (even over periods of up to 150 days without food) [21, 24, 25], actual plasma glucose levels in elasmobranchs are lower than in teleost fish of comparative size and with similar metabolic rates [26]. It has also been found that elasmobranchs have an impressive tolerance of hypoglycaemia, including an ability to cope with a virtual absence of circulating glucose for at least 24 hours, a 75% reduction for at least a week and sub-normal plasma glucose levels for extended periods [22, 27]. There is an obvious paradox associated with an impressive ability to cope with long periods of hypoglycaemia existing in conjunction with the maintenance of apparently stable plasma glucose levels and others have pondered the necessity of central glucose-sensing mechanisms in these species [26].

The chondrichthyan pancreas represents an important model for studies of vertebrate pancreas evolution and function, particularly with reference to glucose homeostasis. However, full analysis of these areas has been hindered by a lack of genetic information and resources - almost the entirety of our current understanding of the chondrichthyan pancreas is based on what might be considered somewhat “old fashioned” (although still vital, important and informative) biological techniques, including descriptive gross anatomy and light and electron microscopy, enzymatic assays typically involving the injection of peptides derived from other (often mammalian) species and immunohistochemistry involving the use of antibodies raised against short mammalian peptide epitopes (see [2, 3, 26, 28] for reviews). There is currently a dearth of data regarding molecular level functions of the chondrichthyan pancreas, including mechanisms of transcriptional and translational control of gene regulation, signaling both in terms of cell-cell communication within the pancreas and in terms of response to neuroendocrine signals and even more basic information such as the sequences of mRNA and protein precursors of previously identified or characterised digestive enzymes and pancreatic peptides.

The lesser spotted catshark (*Scyliorhinus canicula*, often referred to as the lesser spotted dogfish) has recently become the chondrichthyan model of choice for a wide range of genetic, developmental and evolutionary analyses [29] and transcriptomic and genomic sequencing projects are currently underway for this species at Genoscope (www.genoscope.cns.fr). An enigmatic second member of the *Pancreas and duodenal homeobox* (Pdx) gene family (called *Pdx2*) was recently identified in the *S. canicula* pancreas [30] but further studies of the role of this gene or the identification of the presence of additional members of other gene families involved in pancreas development, cell-specification and insulin regulation are impossible without more comprehensive molecular analyses. In order to shed further light on the possible role of the *Pdx2* gene and to go some way to addressing the current dearth of data we set out to determine the pancreas transcriptome of the lesser spotted catshark and to carry out comparative expression analyses with other adult body tissues (liver and brain). These data represent the first large-scale transcriptomic analysis of multiple cartilaginous fish tissues and will be invaluable in understanding the functions of the cartilaginous fish pancreas, as well as shedding light on the evolution of the vertebrate pancreas itself.

## Results

A total of 6,260,398; 32,106,318 and 12,201,682 paired-end sequencing reads were generated for the pancreas, liver and brain respectively and these were pooled to generate a single assembly (Additional file 1) which contained 86,006 contigs (when trimmed to remove all contigs <300bp, which likely represent single pairs of sequencing reads). The tissue distribution of these transcripts was determined by mapping sequencing reads from each tissue to this assembly and abundance values of ≥1 fragments per kilobase per million mapped reads (FPKM) were taken to confirm expression of a particular transcript in each tissue (Table 1). In this way 44,794 contigs were assigned to one or more tissues (Figure 2, Additional files 2-8). All transcripts contained an ORF encoding 20 amino acids or more (Figure 3), of which roughly 4-7% encoded a signal peptide and so are likely to be secreted (Table 1). Typically, between 15-53% of transcripts had a BLAST hit in the RefSeq collection and 12-49% were annotated with GO terms (Table 2). Although low, these figures are broadly comparable with a similar analysis of the white shark heart transcriptome [31], which found matches of 23.5% and 21.5% respectively. In order to provide a broad overview of the assigned gene ontology terms, we carried out a generic GOSlim annotation of the data (Figures 4 and 5), and Fishers exact tests showed that the pancreas was enriched for seven GO terms compared to liver (‘cell’, ‘reproduction’, ‘transcription, DNA-dependent’, ‘embryo development’, ‘growth’, ‘intracellular non-membrane-bounded organelle’ and ‘non-membrane-bounded organelle’) and two terms compared to brain (‘transcription, DNA-dependent’ and ‘cellular amino acid metabolic process’– the full comparative enrichment results are provided in Additional file 9).

**Table 1.** Assembly characteristics. Tissue distribution of transcripts was assigned by read mapping, taking a value of ≥1 fragments per kilobase per million mapped reads (FPKM) as evidence of expression.

| Tissue(s) | Number of<br>contigs<br>>300bp | Contig<br>N50<br>(bp) | Longest<br>contig<br>(bp) | Mean ORF<br>length (bp) | Max ORF<br>length<br>(bp) | Number ORFs<br>with signal<br>peptide |
| --- | --- | --- | --- | --- | --- | --- |
| Pancreas only | 6551 | 429 | 5386 | 177 | 3005 | 257 (3.92%) |
| Brain only | 21241 | 798 | 16085 | 338 | 13751 | 1029 (4.84%) |
| Liver only | 3423 | 676 | 13906 | 271 | 13532 | 220 (6.43%) |
| Pancreas, brain | 3847 | 1479 | 10710 | 482 | 10394 | 210 (5.46%) |
| Pancreas, liver | 924 | 1039 | 5907 | 329 | 4505 | 62 (6.71%) |
| Brain, liver | 2482 | 1329 | 12080 | 481 | 10403 | 149 (6.00%) |
| Pancreas, liver, brain | 6326 | 2257 | 11135 | 705 | 9329 | 356 (5.63%) |

**Table 2.**
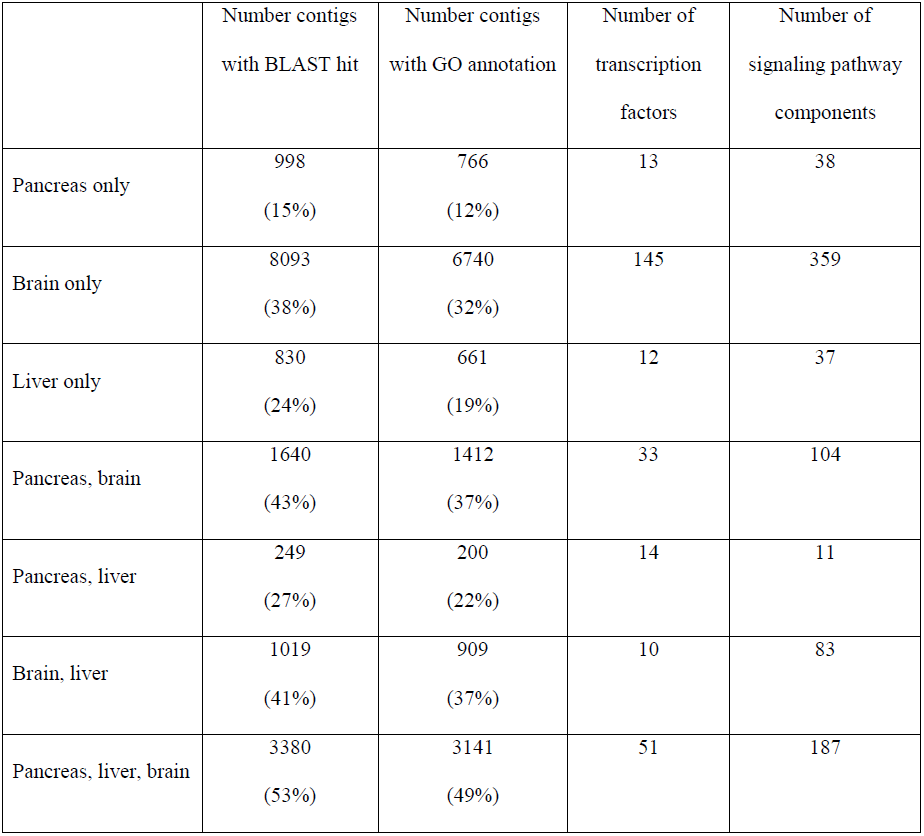
Number of contigs with a BLAST hit in the RefSeq database and Gene Ontology (GO) annotation assigned by BLAST2GO [124, 125]. The number of transcription factors and signaling pathway components in each tissue or tissue combination is also shown.

|  | Number contigs<br>with BLAST hit | Number contigs<br>with GO annotation | Number of<br>transcription<br>factors | Number of<br>signaling pathway<br>components |
| --- | --- | --- | --- | --- |
| Pancreas only | 998<br>(15%) | 766<br>(12%) | 13 | 38 |
| Brain only | 8093<br>(38%) | 6740<br>(32%) | 145 | 359 |
| Liver only | 830<br>(24%) | 661<br>(19%) | 12 | 37 |
| Pancreas, brain | 1640<br>(43%) | 1412<br>(37%) | 33 | 104 |
| Pancreas, liver | 249<br>(27%) | 200<br>(22%) | 14 | 11 |
| Brain, liver | 1019<br>(41%) | 909<br>(37%) | 10 | 83 |
| Pancreas, liver, brain | 3380<br>(53%) | 3141<br>(49%) | 51 | 187 |

**Figure 2.**
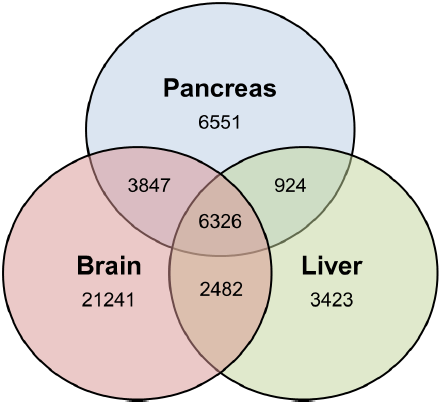
Tissue distribution of transcripts, as determined by mapping the sequencing reads derived from each tissue to a combined, all-tissue assembly. Contig values of ≥1 FPKM (<u>F</u>ragments <u>P</u>er <u>K</u>ilobase of exon per <u>M</u>illion fragments mapped) were taken as evidence for expression.

**Figure 3.**
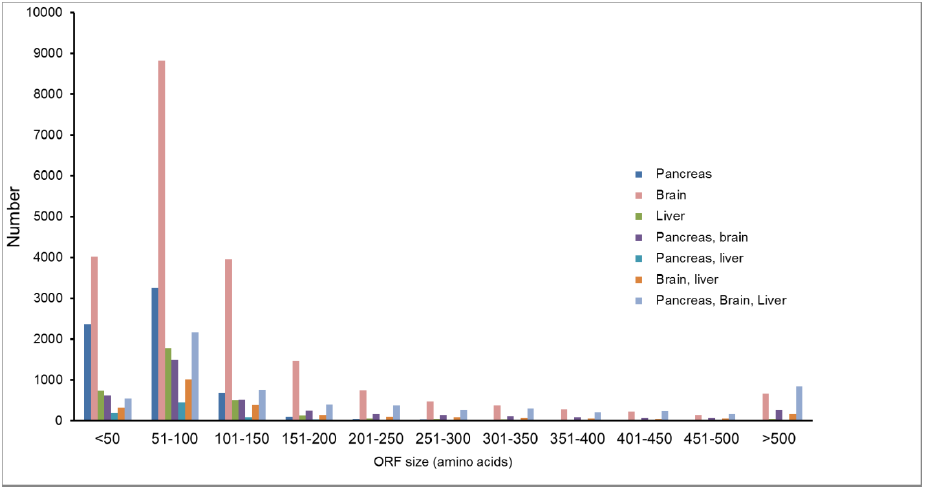
Length and tissue distribution of open reading frames (ORFs) derived from assembled contigs

**Figure 4.**
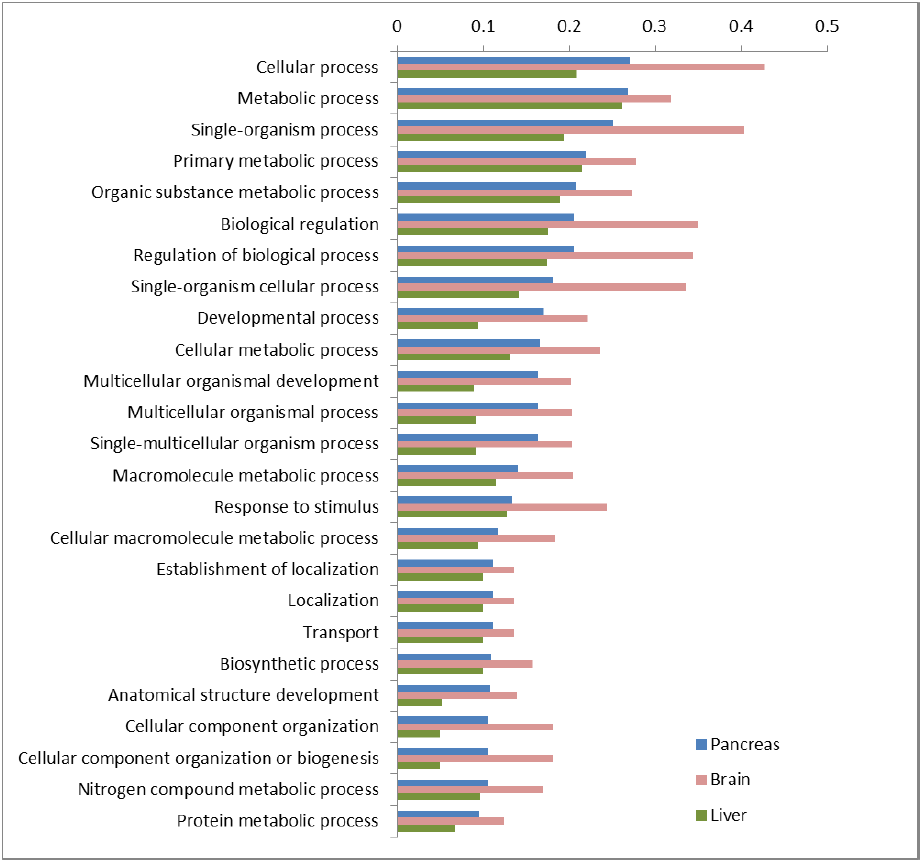
Proportion of transcripts assigned to each of the top 25 gene ontology (GO) slim ‘Biological Process’ terms for catshark pancreas, brain and liver tissue-specific transcripts.

**Figure 5.**
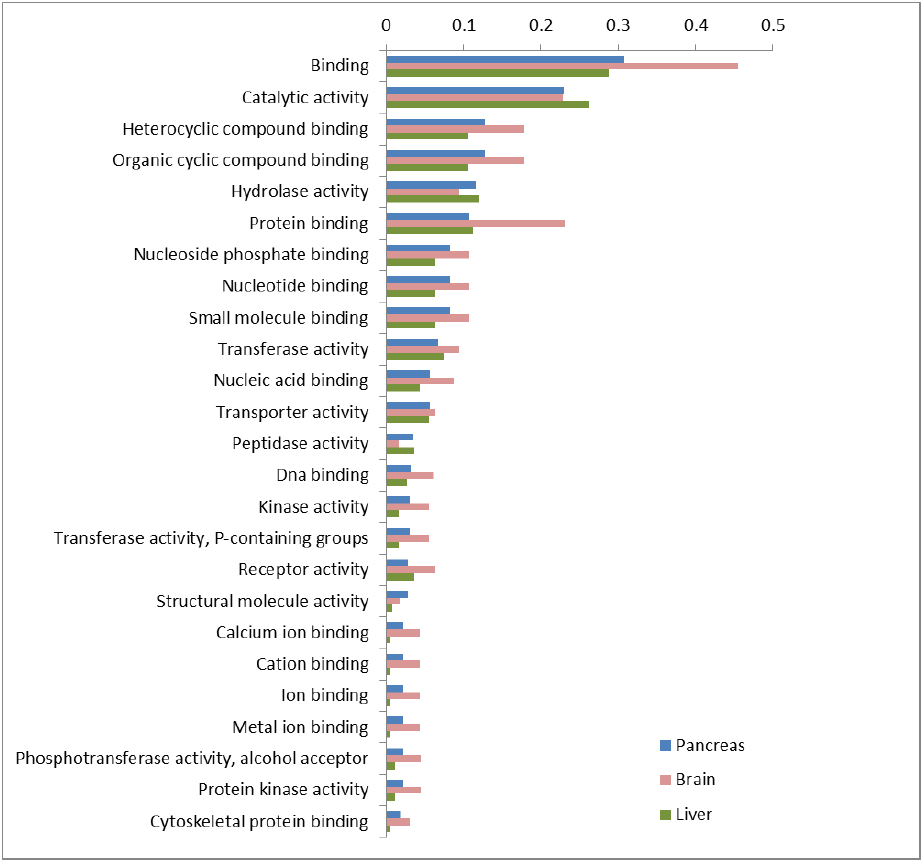
Proportion of transcripts assigned to each of the top 25 gene ontology (GO) slim ‘Molecular Function’ terms for catshark pancreas, brain and liver tissue-specific transcripts.

A more detailed search strategy was carried out for particular categories of genes that would shed light on the similarities or differences of pancreas function in *S. canicula* compared to other vertebrates. The results of these analyses are outlined in the following sections.

### Pancreatic hormones and their receptors

A large amount of immunohistochemical research has putatively identified the presence of a number of pancreatic peptide hormones in cartilaginous fish, including insulin, glucagon, somatostatin and pancreatic polypeptide, and the presence of at least some of these peptides has been confirmed by proteomic studies [11, 12, 15, 32]. Our transcriptomic data confirms the presence of mRNA transcripts encoding preproinsulin, preproglucagon (Figure 6) and preprosomatostatin (corresponding to the *SSa* gene [33]) in the pancreas and preprosomatostatin b and c in both the brain and pancreas, but not pancreatic polypeptide (PP – see next section), gastrin, gastric inhibitory polypeptide (GIP) or secretin. Vasoactive intestinal polypeptide (VIP) is expressed by both pancreas and brain, cholecystokinin by only brain and, as previously suggested [3], ghrelin appears to be absent from the shark pancreas and, indeed, from all three tissues. We find transcripts encoding the insulin receptor (IR) only in brain, the glucagon receptor only in liver and somatostatin receptor 1 (SSTR1) and SSTR5 in brain and both pancreas and brain respectively. Contrary to the findings of Larsson *et al.* [34], we find Neuropeptide Y receptors Y1, Y5 and Y6 in brain only and Y8 in both pancreas and brain, suggesting that the expression of these receptors either varies between chondrichthyan species, or is more dynamic than previously assumed.

**Figure 6.**
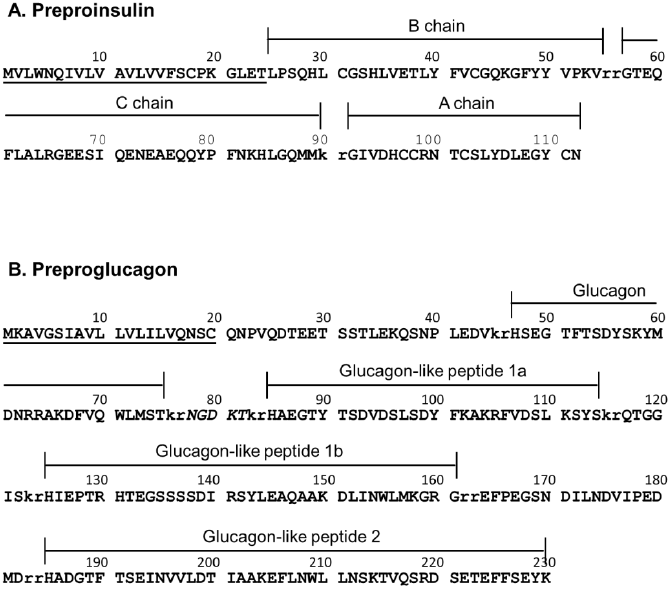
Annotation of the precursor peptides of catshark preproinsulin and preproglucagon. Signal peptides (amino acids 1-24 and 1-20 respectively) are underlined and basic amino acid cleavage sites are lowercase. Glucagon-like peptides (GLP) 1a and 1b are annotated based on similarity to the duplicated GLP1 peptides in the unpublished *Squalus acanthias* and *Hydrolagus colliei* proglucagon sequences available on Genbank (accession numbers AAS57653 and AAS57654). An oxyntomodulin-like peptide has been purified from *H. colliei* and corresponds to amino acids 47-82 in the Catshark preproglucagon.

The presence of PP and γ-cells are key aspects of current schemes for the mode of vertebrate pancreas evolution [1–3, 35]. However, it has been known for some time that PP is tetrapod-specific, produced via duplication of the *Peptide YY* gene sometime prior to the divergence of this lineage [36, 37] and there is therefore a discrepancy between the findings of decades of immunohistochemical research and data from molecular genetic studies and analyses of vertebrate whole genome sequences. We have identified transcripts of two members of the Neuropeptide Y family (which includes Neuropeptide Y (NPY), Peptide YY (PYY) and Pancreatic polypeptide (PP)) in our dataset - a *PYY* gene expressed in pancreas and brain and a *NPY* gene expressed only in brain (Figure 7, both sequences are identical to published catshark sequences for PYY and NPY (accessions P69095 [38], AAB23237 [14]). We therefore suggest that older immunohistochemical studies which claimed to have detected PP+ cells in the cartilaginous fish pancreas may have in fact been relying on antisera that cross-reacted with PYY. A focus on the (often brief) methods sections of several key historical papers revealed that they in fact used the same anti-PP antibody, produced by Ronald Chance at Eli Lily in the 1970’s [16, 39, 40]. It therefore appears that this antibody was detecting PYY in the pancreas of cartilaginous fish and that these initial papers and various subsequent papers have repeatedly been cited until the presence of PP in cartilaginous fish is considered to be established fact. In other cases, the misidentification of sequenced peptides has added to the confusion [38].

**Figure 7.**
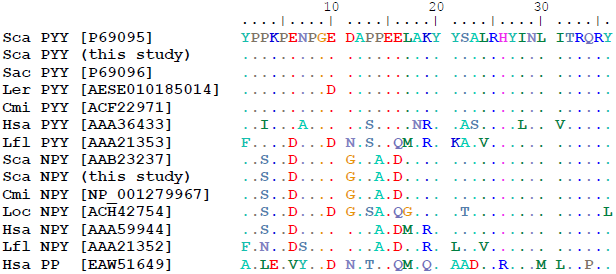
Amino acid alignment of vertebrate Peptide YY (PYY), Neuropeptide Y (NPY) and Pancreatic polypeptide (PP) sequences. Genbank accession numbers are given in square brackets. Sca, Scyliorhinus canicula (lesser spotted catshark); Sac, Squalus acanthias (spiny dogfish); Ler, Leucoraja erinacea (little skate); Cmi, Callorhinchus milii (elephant shark); Hsa, Homo sapiens (human); Lfl, Lampetra planeri (brook lamprey); Loc, Leucoraja ocellata (winter skate).

Our immunohistochemical surveys of the catshark pancreas using high-affinity anti-PP antibodies (Table 1 in Additional file 10) showed varying results. The PP antisera from Sigma weakly stained the catshark pancreas but the staining was completely absorbed with PP, NPY and PYY peptides (Figure 8). While the Millipore anti-PP failed to stain (except on the mouse control pancreas, data not shown), experiments with anti-PYY antibodies detected strong signals, co-localising with insulin but not glucagon or somatostatin (Figure 8). NPY antisera immunoreactivity was detected in the same pattern as PYY (data not shown) and was absorbed with either PYY or NPY peptides (Table 2 in Additional file 10).

**Figure 8.**
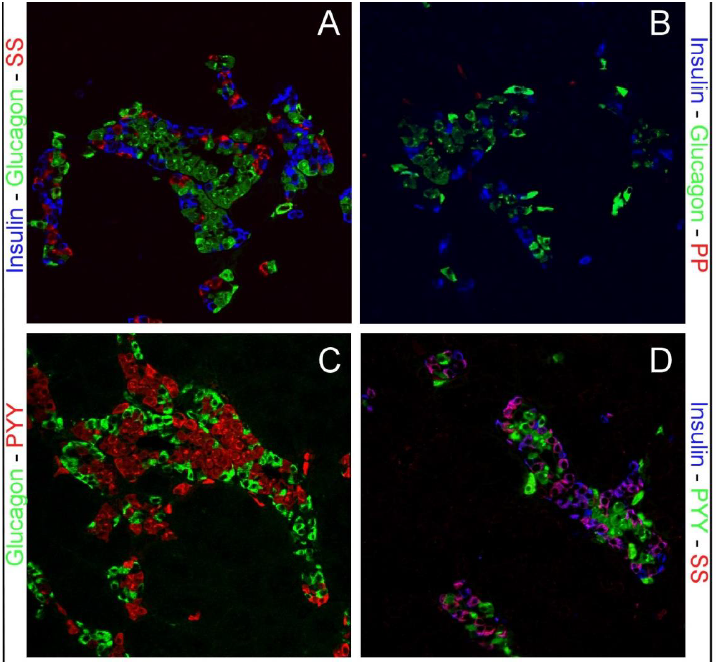
Immunolocalization of pancreatic hormones and pancreatic polypeptide and PYY in catshark pancreas. (A) The distribution of the pancreatic hormones insulin (blue), glucagon (green) and somatostatin (Red) in uniquely shaped islet structures. (B) Pancreatic polypeptide (red) specific antisera fail to stain a specific subset of endocrine cells in the pancreas, while insulin (blue) and glucagon show a normal distribution. (C-D) PYY shows colocalization with most of the insulin immunoreactive cells but not glucagon or somatostatin. All images are 250x magnification

A search of the Blast2GO results for the term ‘Hormone activity’ identifies three other peptide-encoding transcripts expressed in the catshark pancreas: Gastrin-releasing peptide (GRP, in pancreas and brain), which fulfils a variety of roles in the gastrointestinal tract, including the regulation of hormone release and the secretion of pancreatic enzymes [41, 42]; Neuromedin U (NMU, in all three tissues) which is expressed in nerves throughout the gastrointestinal tract [43] and Enkephalin (in pancreas and brain), an endogenous opioid that functions as a neurotransmitter or neuromodulator [44, 45].

### Glucose sensing

*Hexokinase type IV*, more typically called *Glucokinase* (*GK*), is a glucose-phosphorylating enzyme that has been shown to be the key molecule for glucose sensing in mammalian liver and pancreas cells [46] and mutations in *GK* are known to cause Maturity Onset Diabetes of the Young Type II (MODY2) [47, 48]. Somewhat surprisingly, we find that *GK* is expressed in the shark brain and *glucokinase regulatory protein* (*GKRP*) only in liver. We also identified transcripts in all three tissues corresponding to *6-phosphofructo-2-kinase/fructose-2,6-bisphosphatase 2* (*Pfkfb2*), another regulator of glucose metabolism via interaction with GK. Finally we detected transcripts encoding two glucose transporters in all three tissues, namely the solute carrier family 2 genes *Glut1* and *Glut2* and the shark pancreas therefore appears to represent an ancestral state where both types of transporter are expressed in the pancreas, as opposed to the the human pancreas (which relies on *GLUT1*) and that of rodents (which rely more heavily on *Glut2*) [49–51].

### Insulin regulation

The regulation of the insulin gene has, for obvious reasons, been an area of intensive study and a number of key regulators are now known from studies in rodents and humans. Perhaps one of the most important is the *Pancreas and duodenal homeobox 1* (*Pdx1*) gene, also called *Insulin promoter factor 1* (Ipf1), *Islet/duodenum homeobox 1* (*Idx1*) or *Somatostatin transactivating factor 1* (*Stf1*) [52–54] which (in addition to roles in embryonic development and β-cell specification) binds to the TAAT motif-containing A boxes of the mammalian insulin promoter to stimulate transcription [55–58]. Mutations in *PDX1* have been linked to Maturity Onset Diabetes of the Young Type IV (MODY4) [59] and pancreatic agenesis [60, 61]. Cartilaginous fish and coelacanths have previously been shown to have retained an ancient paralogue of *Pdx1*, which we termed *Pdx2* [30, 62] and accordingly we find transcripts of both genes in our pancreas transcriptome dataset. The basic helix-loop-helix (bHLH) transcription factor *NeuroD1* (previously called β2 [63]) has roles in pancreas development and islet formation [64, 65] and mutations in this gene have been linked to Type 2 Diabetes mellitus and MODY6 [66]. *NeuroD1* has also been shown to be important for insulin gene expression [67–69] and the LIM-homeodomain transcription factor *Isl1* acts synergistically with *NeuroD1* and the bHLH transcription factor *E47* to bind to and activate the insulin gene promoter [70–72]. The *hepatocyte nuclear factor 1 alpha* (*Hnf1a*) gene was originally described as a liver-specific transcription actor, responsible for the regulation of a number of genes important for liver function [73–75]. However, it was later found that this gene also has a role in glucose homeostasis via regulation of insulin secretion [76, 77] and that mutations in *Hnf1a* were the cause of MODY3 [78–80]. The *Nkx6.1* homeodomain transcription factor is a potent transcriptional repressor with a key role in β-cell differentiation [81, 82] and has also been shown to suppress the expression of glucagon to maintain β-cell identity, as well as being able to regulate glucose-sensitive insulin secretion [83]. Our discovery of *Pdx1* (and *Pdx2*), *NeuroD1* and its partner *E47*, *Isl1*, *Hnf1a* and *Nkx6.1* transcripts expressed in the catshark pancreas suggests an ancient role for these genes in vertebrate pancreas function and hints at early establishment of the insulin gene regulatory network. Additionally, the presence of transcripts encoding *NeuroD1, e47, Isl1* and *Nkx6.1* in the catshark brain highlights shared ancestry of these tissues in the vertebrate neuroendocrine system. We do not find any transcripts for *MafA*, which has been shown to be a key regulator of glucose-sensitive insulin secretion in humans and rodents [58, 84, 85], although other studies have also had difficulty identifying transcripts of this gene and other pancreas transcription factors in non-PCR based experiments [86, 87], possibly because of the low level of expression of transcription factors in general [88].

### Transcription factors

In addition to the transcription factors involved in insulin regulation discussed above, KEGG orthology (KO) analysis [89–91] has identified 13 transcription factors expressed in just the pancreas (including *Pdx1* and *Pdx2*, *FoxA1* (*Hnf3a*) and *Pancreas-specific transcription factor 1a* (*Ptf1a*)), 14 expressed in pancreas and liver, 33 in pancreas and brain and 51 in all three tissues (Tables 2 and 3).

**Table 3.** Transcription factors and signaling pathway components unique to the lesser spotted catshark pancreas based on available data.

| Pancreas-specific transcription factors | Pancreas-specific signaling components |
| --- | --- |
| Forkhead box protein A1 | 1D-myo-inositol-triphosphate 3-kinase |
| GATA-binding protein 4 | 5-hydroxytryptamine receptor 2 |
| Histone-lysine N-methyltransferase MLL1 | Adenylate cyclase 2 |
| Homeobox protein hoxB5 | Bone morphogenetic protein 2 |
| Homeobox protein hoxB7 | Bone morphogenetic protein 7 |
| Homeobox protein MOX | Cholecystokinin A receptor |
| Pancreas and duodenal homeobox 1 | Collagen, type I/II/III/V/XI/XXIV/XXVII, alpha |
| Pancreas and duodenal homeobox 2 | Collagen, type IV, alpha |
| Krueppel-like factor 5 | Collagen, type VI, alpha |
| Nuclear receptor subfamily 0 group B 1 | Cysteinyl leukotriene receptor 1 |
| Nuclear receptor subfamily 0 group B 1 | Epidermal growth factor receptor |
| Pancreas-specific transcription factor 1a | Fibroblast growth factor |
| Transcriptional enhancer factor | FMS-like tyrosine kinase 1 |
| Zinc finger protein GLI3 | Frizzled 9/10 |
|  | Glutamine synthetase |
|  | Inositol 1,4,5-triphosphate receptor type 1 |
|  | Inositol 1,4,5-triphosphate receptor type 2 |
|  | Inositol 1,4,5-triphosphate receptor type 3 |
|  | Insulin |
|  | Integrin alpha 1 |
|  | Integrin alpha 2 |
|  | Integrin beta 6 |
|  | Interleukin 10 receptor beta |
|  | Janus kinase 2 |
|  | Laminin, alpha 3/5 |
|  | Laminin, beta 4 |
|  | Leucine-rich repeats and death domain-containing protein |
|  | Mitogen-activated protein kinase kinase kinase kinase 4 |
|  | Nucleoprotein TPR |
|  | Protein crumbs |
|  | Receptor-interacting serine/threonine-protein kinase 1 |
|  | Secretory phospholipase A2 |
|  | Suppressor of cytokine signaling 1 |
|  | TGF-beta receptor type-2 |
|  | Transcriptional enhancer factor |
|  | Transferrin |
|  | Vascular endothelial growth factor C/D |
|  | Zinc finger protein GLI3 |

In a survey of the expression of 790 human DNA-binding transcription factors, Kong et al. [88] identified 80 with expression restricted to the fetal pancreas, 32 restricted to the adult pancreas and 18 shared by both. Of the 31 adult-specific genes, we find evidence that 6 are also expressed in the adult catshark pancreas, although this number increases to 15 if members of the same gene family are considered (the possibility of divergent resolution of gene duplicates following the whole genome duplications [92] in early vertebrate ancestry must be considered). Since transcription factors are known to be expressed at low levels in cells (less than 20 copies per human adult cell [88]) it is likely that our figure is an underestimate and a more comprehensive survey of candidate transcription factor expression in this species is needed.

### Signalling

Our KEGG orthology analysis identified 38 transcripts involved in signal transduction that are expressed only in the catshark pancreas, 11 in both pancreas and liver, 104 in pancreas and brain and 187 in all three tissues (Tables 2 and 3). Among these are representatives of the major vertebrate signalling pathways, including ligands and receptors for Fgf, Wnt, Notch, Vegf, Tgfβ and Pdgf. Members of all of these pathways have previously been identified in the human pancreas transcriptome [87].

### Homeobox gene diversity

Homeobox genes are a group of transcription factors that encode a 60 amino acid DNA-binding homeodomain and that are involved in a wide variety of gene regulatory events in embryonic and adult tissues. A number of homeobox genes are known to be expressed during endodermal regionalisation and pancreas development, including *Islet 1* and *2* (*Isl1*, *Isl2*), *Pancreatic and duodenal homeobox 1* (*Pdx1*), *Nkx6.1*, *Nkx2.2*, *Pituitary homeobox 2* (*Pitx2*), *Motor neuron and pancreas homeobox 1* (*Mnx1*), *Onecut homeobox 1* (*Onecut/Hnf6*) and *Paired box genes 4* and *6* (*Pax4*, *Pax6*) [93, 94]. Some older studies have detected a variety of homeobox genes in mammalian pancreas cell lines, including *Cdx4*, *Hox1.4* (*HoxA4*), *Chox7* (*Gbx1*), *Hox2.6* (*HoxB4*), *Cdx3* (*Cdx2*), *Cdx1*, *Hox4.3* (*HoxD8*), *Hox1.11* (*HoxA2*), *Hox4a* (*HoxD3*), *Hox1.3* (*HoxA5*) in the somatostatin-producing rat insulinoma cell line RIN1027-B2 [53] and *Isl1*, *Lmx2*, *Alx3*, *HoxA4*, *HoxA13*, *Ipf1* (*Pdx1*), *Nkx2.2*, *Nkx6.1*, *En2* and *Vdx* in a hamster insulinoma cell line [95]. More recently, microarray and RNA-seq studies have identified a much larger number of homeobox genes expressed in the pancreas and especially the β-cell, with over 60 different homeobox genes identified by Kutlu et al. [87]. We used the homeodomain sequences of all human homeobox genes from HomeoDB [96, 97] and all vertebrate homeobox gene sequences from Pfam [98] as BLAST queries against our catshark transcriptome data and identified 11 different homeobox genes expressed in the pancreas, including five in just pancreas (*HoxB5, HoxB7, Mox1, Pdx1, Pdx2*), two in pancreas and liver (*Hlx, Hhex*), three in pancreas and brain (*Arx, Zfhx3, Zfhx4*) and one in all three tissues (*Cut-like 2*). These include genes known to be restricted to, or highly expressed in, β-cells (*Pdx1*), α-cells (*Arx*) and acinar cell types (*Cut-like 2*) [86].

### Digestion

In addition to its endocrine roles, the pancreas is also an important exocrine organ, fulfilling key functions in the digestion of proteins, lipids and carbohydrates. In the carnivorous elasmobranchs protein and lipids are the main energy sources [99] and it has been shown that ketone bodies and amino acids are the main oxidative fuel source for muscles and several other tissues, in preference to fatty acids [24, 28, 99].

Carbohydrates are thought to be utilised as oxidative fuels in elasmobranch heart muscle, as well as brain, red muscle and rectal gland [28, 100, 101]. It is therefore perhaps reasonable to assume that proteases and lipases are the most significant digestive enzymes produced by the elasmobranch pancreas and indeed this appears to be the case. Some form of chymotrypsinogen and trypsinogen have long been known to be produced by the elasmobranch pancreas, as has carboxypeptidase B, although these enzymes have not been fully characterised or isolated and sequenced [102–105]. We find transcripts of *Elastase 2a* and *3b*, *Chymotrypsinogen b1* (*Ctrb1*), *Chymotrypsin-like* (*Ctrl*), *Chymotrypsin-like elastase family, member 1* (*Cela1*) and *Chymotrypsin-like elastase family, member 3B* (*Cela3b*), *Trypsin 1, 2* and *3*, as well as the digestive carboxypepetidases (A1, A2, B1) and those involved in activation and processing of other proteins, such as *carboxypeptidase B2, D* and *E* [106, 107].

Some form of triacylglycerol lipase activity has previously been detected using crude enzyme preparations from the pancreas of skate (*Raja* (now *Amblyraja*) *radiata*) [108] and Leopard shark, *Triakis semifasciata* [109]. However, we find no evidence of Pancreatic liapse in the catshark pancreas transcriptome and instead find only Pancreatic lipase-related proteins 1 (Pnliprp1) and 2 (Pnliprp2). It is likely therefore that the triacylglycerol lipase activity found previously is a result of the action of Carboxyl ester lipase (CEL or bile salt-stimulated lipase) [110]. We also find colipase, in agreement with earlier studies of a range of cartilaginous fish and other basal vertebrates [111–113] and Hepatic and Hormone-sensitive lipases. Several lipid transporting apolipoproteins are also expressed by the catshark pancreas, including apolipoproteins A-IV, E, M and O. Finally, we have identified transcripts of genes involved in the digestion of carbohydrates (Pancreatic alpha amylase) and nucleic acids (deoxyribonuclease I and various ribonucleases).

### Microsatellites

It has recently been suggested [31] that a high frequency of dinucleotide simple sequence repeats (SSRs, microsatellites) is a general feature of shark genomes. We find 6,843 transcripts containing one or more di-, tri- or tetranucleotide microsatellites of five perfect repeats or more in our catshark data, with 482 of these only in pancreas, 3,083 only in brain and 473 only in liver (Table 4). In accordance with previous suggestions [31] we find dinucleotide repeats to be the most common type of SSR in both coding and non-coding regions of catshark transcripts.

**Table 4.** Predicted di-, tri- and tetranucleotide simple sequence repeats (microsatellites) in each catshark tissue or combination or tissues. Results are shown for both full transcripts and predicted open reading frames (ORFs) and in both cases dinucleotide repeats are the most common.

|  | Transcript |  |  | Predicted ORFs |  |  |
| --- | --- | --- | --- | --- | --- | --- |
|  | Di- | Tri- | Tetra- | Di- | Tri- | Tetra- |
| Pancreas only | 456<br>(6.96%) | 23<br>(0.35%) | 3<br>(0.05%) | 60<br>(0.92%) | 6<br>(0.09%) | 0<br>(0%) |
| Brain only | 2808<br>(13.22%) | 268<br>(1.26%) | 7<br>(0.03%) | 829<br>(3.90%) | 147<br>(0.69%) | 1<br>(<0.01%) |
| Liver only | 446<br>(13.03%) | 24<br>(0.70%) | 3<br>(0.09%) | 168<br>(4.90%) | 12<br>(0.35%) | 1<br>(0.03%) |
| Pancreas +<br>Brain | 481<br>(12.50%) | 43<br>(1.12%) | 2<br>(0.05%) | 72<br>(1.87%) | 15<br>(0.39%) | 2<br>(0.05%) |
| Pancreas +<br>Liver | 105<br>(11.36%) | 5<br>(0.54%) | 1<br>(0.11%) | 44<br>(4.76%) | 3<br>(0.32%) | 1<br>(0.11%) |
| Brain + Liver | 702<br>(28.28%) | 61<br>(2.46%) | 3<br>(0.12%) | 106<br>(11.47%) | 21<br>(2.27%) | 0<br>(0%) |
| Pancreas +<br>Liver + Brain | 1240<br>(19.60%) | 158<br>(2.50) | 4<br>(0.06%) | 308<br>(4.87%) | 108<br>(1.71%) | 2<br>(0.03%) |

## Discussion

Our analysis of the catshark pancreas transcriptome reveals the presence of genes known to be involved in glucose sensing and regulation of the insulin gene in other vertebrates and illustrates that functional conservation of these aspects of the vertebrate pancreas is reflected at the molecular-level. We therefore propose that these molecular-level mechanisms are a common feature of jawed vertebrates and that this lends support to the theory that the evolution of blood-glucose sensing and regulatory mechanisms may have facilitated the evolution of the complex glucose-dependent brain of vertebrates [7–9]. We further suggest that the early evolution and fixation of these mechanisms has imposed evolutionary constraints on glucose sensing and insulin regulation in vertebrates, including in cartilaginous fish, even in the face of their ability to tolerate extended periods of hypoglycaemia and likely relaxed requirements for these processes.

We find that the catshark pancreas produces at least eight peptide hormones (insulin; glucagon; somatostatin; peptide YY; gastrin-releasing peptide, neuromedin U, encephalin and vasoactive intestinal polypeptide, Table 5) and expresses a wide variety of genes involved in digestion, especially the digestion of proteins and lipids. The catshark pancreas therefore clearly has the features of a distinct pancreatic gland with both endocrine and exocrine functions and as such will be of great use in reconstructing the characteristics of the earliest vertebrate pancreas. The similarity in gene expression between the catshark and other vertebrates with respect to hormones, digestive enzymes, transcription factors and signaling pathways again provides support to the theory that there was a single, early origin of the pancreas at the base of the jawed vertebrate radiation. The overlap in peptides produced by the catshark pancreas and brain (Table 5) is a reflection of the shared ancestry of these tissues within the vertebrate neuroendocrine system [114].

**Table 5.** Peptide diversity of the catshark pancreas, brain and liver. Our comprehensive transcriptomic survey of the lesser spotted catshark pancreas highlights the disparity in the estimation of peptide diversity in early vertebrates as previously suggested by immunohistochemical (IHC) studies and highlights the similarity of pancreas and brain peptide complements.

| **Peptide** | **Pancreas**<br>**IHC** | **Pancreas**<br>**transcriptome** | **Brain**<br>**transcriptome** | **Liver**<br>**transcriptome** |
| --- | --- | --- | --- | --- |
| Insulin | + | + | - | - |
| Glucagon | + | + | - | - |
| Somatostatin | + | + | + | - |
| Pancreatic polypeptide | + | - | - | - |
| Peptide YY | + | + | + | - |
| Gastrin-releasing peptide | + | + | + | - |
| Neuromedin U | + | + | + | + |
| Enkephalin | + | + | + | - |
| Cholecystokinin | + | - | + | - |
| Gastrin | + | - | - | - |
| Vasoactive intestinal polypeptide | + | + | + | - |
| Gastric inhibitory polypeptide | + | - | - | - |
| Secretin | + | - | - | - |

Based on its co-localisation with insulin-, glucagon-, somatostatin- and PP-cells during mouse development, it has previously been suggested that a PYY+ cell may constitute a common progenitor of the major islet cell types [115]. Recent lineage tracing experiments have demonstrated that PYY+ cells give rise to islet ∂ and PP cells and approximately 40% of pancreatic α and at least some β cells arose from peptide YY+ cells [116]. Most β cells and the majority of α cells are therefore not descendants of the peptide YY+/glucagon+/insulin+ cells that first appear during early pancreas ontogeny. The co-localisation of PYY with insulin in the adult shark pancreas illustrates the diversity of mechanisms that exist in vertebrate pancreas development and function and demonstrates the utility of “non-model” species to study these processes. The catshark PYY+ cells will therefore provide important insights into the evolution of the vertebrate pancreas, and especially progenitors of α, β, δ and γ-cells.

Our experiments make clear that much of the previous work on the presence or absence of peptides in basal vertebrate lineages may be suspect, with many false-positive signals resulting from cross-reacting antisera. Previous schemes of pancreas evolution based on these and similar data, which posited the restriction of various hormones to the alimentary canal (similar to the situation in protochordates such as amphioxus), the accumulation of these into a two-or three peptide islet organ in jawless fish and finally the “classic four-hormone islet tissue” of cartilaginous fish and other vertebrates [2] are therefore incorrect. In fact, it appears that the three hormone (glucagon, insulin, somatostatin) islet organ was established early in vertebrate evolution and remains today in the adult (but not larval) lamprey, cartilaginous fish and actinopterygian (ray-finned) fish, and that it is only in the sarcopterygian (lobe-finned fish) lineage that a four hormone (the above, plus PP) pancreas was formed.

Our analysis of homeobox gene expression reveals a surprising level of variation between the genes known to be expressed in the catshark pancreas, human islets [87] and rat [53] and hamster [95] cell lines. It therefore seems likely that this particular class of transcription factors is extremely variable with respect to their spatial or temporal expression pattern in the vertebrate pancreas (or more likely both) and this is perhaps not too surprising given the variety of roles carried out by the pancreas in response to feeding, digestion and the regulation of blood glucose. As expected we have identified transcripts of both *Pdx1* and *Pdx2* in the catshark pancreas, although we do not find any evidence for the presence of additional duplicates of other genes encoding proteins known to interact with PDX1 in other species. It therefore seems unlikely that the maintenance of paralogous *Pdx2* genes in some vertebrate lineages reflects a wider conservation of duplicated gene regulatory networks produced as a result of whole genome duplication events early in vertebrate evolution. Comparison of the amino acid sequences of PDX1 and PDX2 across vertebrates shows conservation of the Pbx-interacting motif, DNA-binding domain and nuclear localisation signal but not of known transactivation domains and the PCIF1-interaction domain [117–119] (Figure 9). The functions of the *Pdx2* gene and the reasons for its retention in some species and independent loss in others (ray-finned fish and tetrapods) remain unknown.

**Figure 9.**
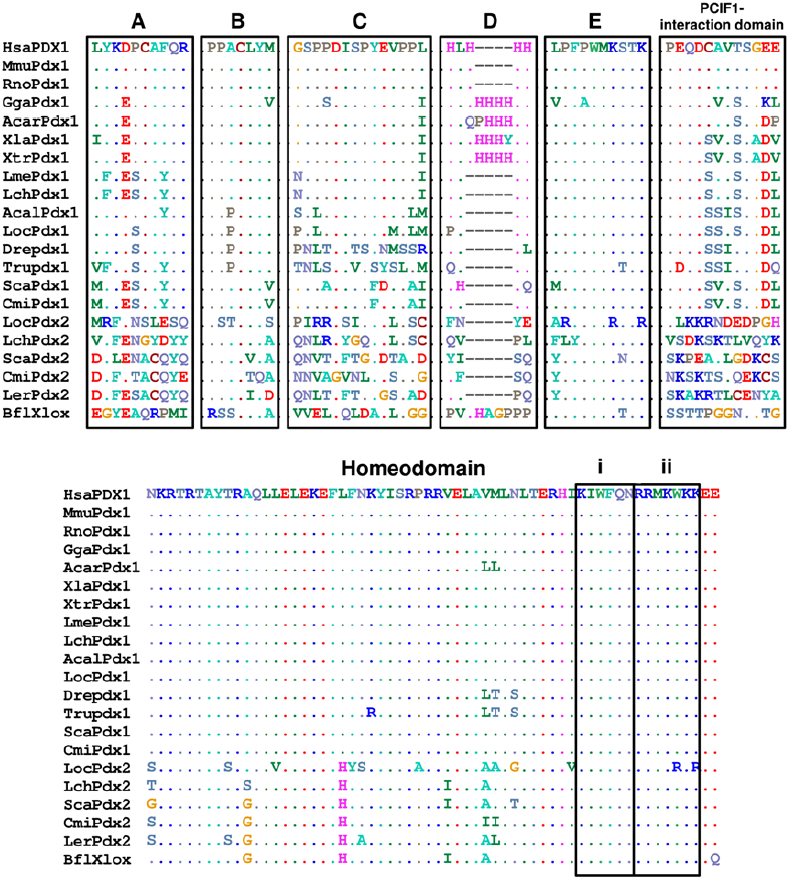
Protein domains in vertebrate PDX1 and PDX2. Transactivation domains A-E [119], PCIF1-interaction domains [118], homeodomains, DNA-binding domains (i) and nuclear localisation signals (ii) are highlighted. Domain E contains the PBX-interacting hexapeptide motif [117]. There is very little conservation of amino acid sequence between the paralogous PDX1 and PDX2 suggesting that they carry out distinct functions within the pancreas, although clearly both are localised to the nucleus, bind DNA and interact with PBX proteins. Hsa, human (*Homo sapiens*); Mmu, mouse (*Mus musculus*); Rno, rat (*Rattus norvegicus*); Gga, chicken (*Gallus gallus*); Acar, Anole lizard (*Anolis carolinensis*); Xla, *Xenopus laevis*; Xtr, *Xenopus tropicalis*; Lme, Indonesian coelacanth (*Latimeria menadoensis*); Lch, African coelacanth (*Latimeria chalumnae*); Acal, Bowfin (*Amia calva*); Loc, Spotted gar (*Lepisosteus oculatus*); Dre, zebrafish (*Danio rerio*); Tru, fugu (*Takifugu rubripes*); Ola, medaka (*Oryzias latipes*); Gac, stickleback; Tni, Green spotted puffer (*Tetraodon nigroviridis*); Sca, lesser spotted catshark (*Scyliorhinus canicula*); Ler, little skate (*Leucoraja erinacea*); Cmi, elephant shark (*Callorhinchus milii*); Bfl, amphioxus (*Branchiostoma floridae*)

With the availability of whole genome sequence information for a greater number of taxa and improved coverage of vertebrate pancreas transcriptomes a larger amount of data than ever before is now becoming available. These data, together with an appreciation that early vertebrate evolution was characterised by extensive genetic, developmental and morphological innovation facilitated by multiple whole genome duplications [120, 121] will better enable us to reconstruct pancreas evolution. As an example, we propose that the creation of the paralogous NPY and PYY during these duplications [34] facilitated the separation of the neuronal and gastroenteropancreatic (GEP) endocrine systems. We further suggest that the availability of additional copies of developmentally-important genes produced during the same duplication events [121] enabled the remodelling of the developing gut and the formation of a distinct pancreas with both endocrine and exocrine functions.

## Conclusions

We have generated a multi-tissue transcriptomic resource for an up and coming model organism, the lesser spotted catshark, *Scyliorhinus canicula*. Somewhat surprisingly we find few transcripts in common between the liver and pancreas, despite their relatively similar roles and shared developmental history as endodermal neighbors. The higher number of transcripts in common between brain and pancreas may provide evidence in support of the co-opting of neuronal programs by at least some pancreatic cells during vertebrate evolution [114, 122], although further comparative analyses are needed in this area. The similarity between the catshark pancreas transcriptome and those of various mammals with respect to insulin regulation, transcriptional and signaling machinery and peptide hormones and their receptors supports the single, early origin of a distinct pancreatic gland in vertebrates, although it seems likely that the peptide diversity of the early vertebrate pancreas may have been overestimated by older, immunohistochemical studies. The cartilaginous fish have a three peptide (insulin, glucagon and somatostatin) pancreas and the four peptide system seen in actinopterygian (ray-finned) and sarcopterygian (lobe-finned) fish and tetrapods is a later evolutionary innovation. The retention of the Pdx2 gene in cartilaginous fish does not apparently reflect a wider retention of duplicated members of pancreas gene regulatory networks and the possible function(s) of this gene remains enigmatic. Our data, together with available or in progress transcriptomic and genomic resources for this and other chondrichthyan species will greatly facilitate comparative studies of elasmobranch, chondrichthyan and vertebrate evolution, particularly with reference to energy metabolism and the maintenance of stable blood glucose levels.

## Methods

### RNA-Seq and sequence analysis

Experimental methods involving animals followed institutional and national guidelines and were approved by the Bangor University Ethical Review Committee. Total RNA was extracted from freshly-dissected pancreas, liver and brain of two adult male and female catsharks approximately 24 hours post feeding. Pancreas samples were sequenced with using 2 × 250bp paired-end reads on the Illumina MiSeq platform at the Centre for Genomic Research (CGR) at the University of Liverpool. Brain and liver samples were sequenced using 2x 150bp paired-end reads on the Illumina HiSeq 2000 platform at the Institute of Biological, Environmental & Rural Sciences (IBERS) at Aberystwyth University. Sequencing reads from the three tissues were assembled into a global tissue assembly using Trinity [123] with the jellyfish K-mer counting method. Tissue distribution of transcripts was assessed by mapping sequencing reads from each tissue to this global assembly, with an FPKM (fragments per kilobase per million mapped reads) value of >1 taken as confirmation of expression. Transcript annotation and assignment of gene ontology (GO) terms was performed using BLAST2GO [124, 125], the KEGG Automatic Annotation Server (KAAS [90]) and by local BLAST using BLAST+ v2.2.27 [126].

### Immunohistochemistry

Male catsharks were euthanized according to a Schedule 1 method and the pancreas removed and fixed in 4% paraformaldehyde/PBS overnight at 4°C. The fixed pancreas was then rinsed several times in PBS, dehydrated through a graded ethanol series and stored in 100% ethanol. 5µm sections of paraffin embedded catshark pancreas were cut and mounted on glass slides. Slides were microwave treated in Tris-EGTA (TEG) buffer ph9.0 and allowed to cool for 30mins. Slides were rinsed in PBS and blocked in normal donkey serum and TNB buffer (Perkin Elmer) for 30mins. Primary antisera were added over night at room temperature and the next day the slides were rinsed 3 × 5min each in PBS and specific cross absorbed donkey anti-mouse, rabbit, or guinea pig secondary antisera (Jackson Immunoresearch) were added for 30mins. The slides were rinsed in PBS and mounted. Details of antisera are given in Table 1 in Additional file 10. All pictures were taken on a Zeiss Meta510 confocal microscope.

### Antibody Absorption

In order to test their specificity against the Pancreatic polypeptide family, the antisera were incubated overnight at 4°C with 10µg of either pancreatic polypeptide (Sigma), Neuropeptide Y (Bachem) or peptide YY (in-house synthesis) or no peptide. The next day the antisera were added to the slides and the staining was performed as above. The staining intensity was compared to the no peptide control and given a rating of 1-3 (+, ++, +++). The results are shown in Table 2 in Additional file 10.

## Authors’ contributions

JFM devised the study and drafted the manuscript; JFM, ADH MJH and MTS carried out RNA-Seq experiments and data analysis, RSH carried out immunohistochemical experiments. All authors read and approved the final manuscript.

## Acknowledgements

This research was supported by a Diabetes UK Small Grant to JFM and donations by the Llandudno and District Diabetes Group. We thank Ashley Tweedale and Gavan Cooke for assistance with collection and maintenance of catsharks and Charity MK McGrae for assistance with immunohistochemistry and imaging. JFM, MJH and MTS are supported by the Biosciences, Environment and Agriculture Alliance (BEAA) between Bangor University and Aberystwyth University and ADH is funded by a Bangor University 125^th^ Anniversary Studentship.

## Additional files

Additional file 1: Combined tissue assembly, trimmed to remove contigs <300bp

Additional file 2: Pancreas-specific transcripts present at ≥1 FPKM, trimmed to remove contigs <300bp

Additional file 3: Brain-specific transcripts present at ≥1 FPKM, trimmed to remove contigs <300bp

Additional file 4: Liver-specific transcripts present at ≥1 FPKM, trimmed to remove contigs <300bp

Additional file 5: Pancreas/brain transcripts present at ≥1 FPKM, trimmed to remove contigs <300bp

Additional file 6: Pancreas/liver transcripts present at ≥1 FPKM, trimmed to remove contigs <300bp

Additional file 7: Brain/liver transcripts present at ≥1 FPKM, trimmed to remove contigs <300bp

Additional file 8: Pancreas/brain/liver transcripts present at ≥1 FPKM, trimmed to remove contigs <300bp

Additional file 9: Gene ontology (GO) enrichment results for pairwise tissue comparisons

Additional file 10: Antibody table and peptide absorption results

## Additional file 10

**Additional table 1.** Table of antisera used in immunohistochemical surveys of the catshark pancreas. PYY, peptide YY; NPY, neuropeptide Y: PP, pancreatic polypeptide

| Antigen | Species | Dilution | Source |
| --- | --- | --- | --- |
| PYY | Rabbit | 1:500 | Sigma |
| PYY | Rabbit | 1:500 | AbCam, UK |
| NPY | Rabbit | 1:1000 | AbCam,UK |
| Insulin | Mouse | 1:75 | Novo Nordisk A/S, Denmark |
| Insulin | Guinea Pig | 1:200 | DAKO, Glostrup, Denmark |
| Glucagon | Mouse | 1:50 | Novo Nordisk A/S, Denmark |
| Glucagon | Rabbit | 1:500 | DAKO, Glostrup, Denmark |
| PP | Rabbit | 1:500 | Sigma |
| PP | Guinea Pig | 1:500 | Linco/Millipore,, |
| Somatostatin | Mouse | 1:100 | Novo Nordisk A/S, Denmark |

**Additional table 2.**
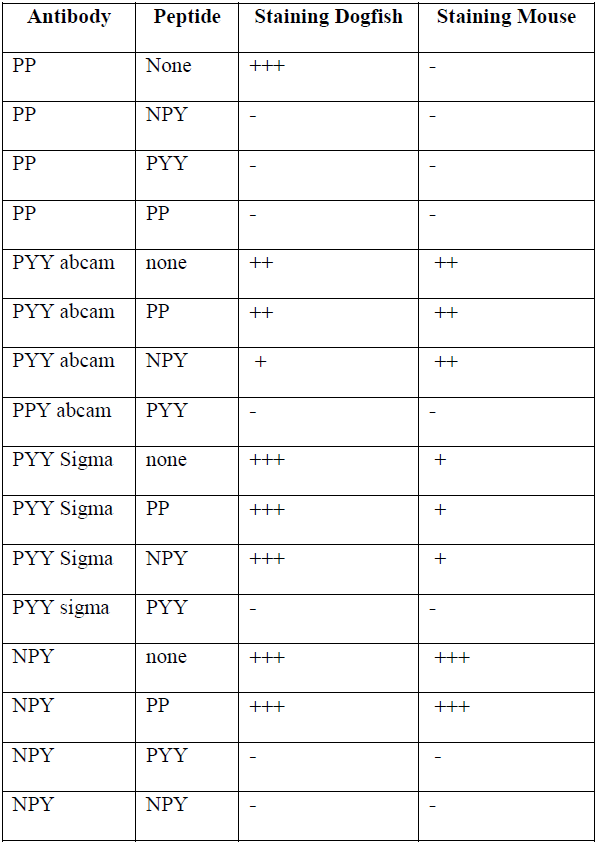
Absorption of Pancreatic Polypeptide Family antisera. Staining is characterised from strong (+++) to weak (+) or absent (-). PYY, peptide YY; NPY, neuropeptide Y: PP, pancreatic polypeptide.

